# tagPAINT: covalent labelling of genetically encoded protein tags for DNA-PAINT imaging

**DOI:** 10.1101/604462

**Authors:** Daniel J. Nieves, Geva Hilzenrat, Jason Tran, Zhengmin Yang, Hugh H. MacRae, Matthew A. B. Baker, J Justin Gooding, Katharina Gaus

**Affiliations:** EMBL Australia Node in Single Molecule Science, School of Medical Sciences, University of New South Wales, Sydney, Australia; ARC Centre of Excellence in Advanced Molecular Imaging, University of New South Wales, Sydney, Australia; Commonwealth Scientific and Industrial Research Organisation (CSIRO), Manufacturing, Clayton, VIC 3168, Australia; School of Biotechnology and Biomolecular Science, University of New South Wales, Sydney, NSW 2052, Australia; School of Chemistry, Australian Centre for NanoMedicine and the ARC Centre of Excellence in Convergent Bio-Nano Science and Technology, University of New South Wales, Sydney, Australia

## Abstract

Recently, DNA-PAINT single molecule localisation microscopy (SMLM) has shown great promise for quantitative imaging. However, labelling strategies so far have relied on approaches that are multivalent or affinity-based. Here, we demonstrate tagPAINT - the covalent labelling of expressed protein tags (SNAP tag and Halo tag) with single DNA docking strands for single molecule localisation microscopy via DNA-PAINT. We utilised tagPAINT for T-cell receptor signalling proteins at the immune synapse as a proof of principle.

## 1. Introduction

An important challenge for single molecule localization microscopy for quantitative measurements is control over the stoichiometry of the label to the molecule of interest. Lack of such control can confound or alter the observed biological behaviour and states of single molecules, complexes and structures due to suboptimal labelling [1, 2], crosslinking due to multivalency, or label-exchange [3]. To address these challenges approaches have been developed that allow a chemically versatile stoichiometric covalent linkage to be formed between biological molecule and fluorescent probes, in structurally defined positions [4, 5]. For example, stable and stoichiometric coupling of fluorescent labels to proteins of interest has been achieved through genetically encoded affinity tags [6, 7], non-canonical amino acid (ncAA) labelling [8, 9], and orthogonal chemistry [10, 11]. Such approaches have then been used to observe proteins at the single molecule level [10, 12].

Recently, Jungmann *et al.* demonstrated a SMLM approach using the binding/unbinding of short fluorescently conjugated DNA probes to antibodies labelled with complementary target strands, known as DNA PAINT [13–16]. This approach was extended to determine the number of proteins/targets present in sub-diffraction structures, termed qPAINT [17]. This method abrogates the uncertainty associated with the stochastic nature of fluorophore blinking and exploits *a priori* knowledge of the binding/unbinding behaviour of the probes. With this approach good agreement was achieved between the theoretical binding/unbinding rate and the observed number of proteins. However, this approach relied on the use of probes labelled with multiple DNA target strands. Thus, multivalent interactions between proteins, and multiple target strands per protein, and incomplete labelling are still challenges that need to be addressed. More recently approaches to address this aim to minimise the linkage error, and include ncAA incorporation [8], affimers [18], fluorescent protein nanobodies [19, 20], and SOMAmers [21] which all allow 1:1 functionalization. However, although SOMAmers spend a long time bound, they still rely on a non-covalent interaction, and can potentially dissociate during prolonged acquisistion times. Similarly, affimers are non-covalent, and sometimes require post-fixation, which may lead to off-target labelling. Also these reagents are only available for a few protein targets to date. Finally, while ncAA incorporation does allow a covalent stoichiometric linkage, it suffers from low expression and efficiency for labelling. New approaches which allow covalent and stoichiometric labelling of a protein of interest, while maintaining a low linkage error, would thus allow robust counting of protein numbers within cell, and thus full sampling of the heterogeneity therein [16].

At present there are a variety of methods to label proteins of interest covalently. One approach is incorporation of an enzymatically active tag, such as SNAP/CLIP tag [22] and Halo Tag [23] technology. The Halo tag makes use of a chemical reaction orthogonal to eukaryotes, *i.e.*, the dehalogenation of haloalkane ligands, thus, leading to highly specific covalent labelling of the tag, and therefore protein [23], in both live and fixed cells. Haloalkanes can be modified to bear fluorescent labels, and has been demonstrated before for single molecule localisation microscopy (SMLM) in live cells using ATTO dye modified ligands [10]. Similarly, SNAP tag, a mutant of DNA repair protein O^6^-alkylguanine-DNA alkyltransferase, can be covalently modified using O^6^-benzylguanine substrates (BG), and has also been demonstrated to be suitable for SMLM imaging [24]. Combining such tagging systems with DNA-PAINT imaging opens up the possibility for robust quantitative imaging of proteins within cells. This is of particular interest as both tagging systems react covalently with a single ligand, thus, the stoichiometry is 1ligand:1 tag, whilst also reducing the linkage error (size of tags c.a. < 5 nm). The labelling of these proteins with DNA oligonucleotides has been demonstrated previously [25, 26], thus, the potential for utilising them for DNA-PAINT imaging is attractive.

Here, SNAP tag and Halo Tag technologies are exploited to allow stoichiometric labelling of single proteins with a DNA PAINT target strand *(i.e.*, 1 protein: 1 target strand), using DNA-modified ligands to the tags. The specificity of the approach is demonstrated by targeting T-cell signalling proteins, CD3ζ and LAT, bearing the tags at the immune synapse in knockout cell lines. The potential of dual-channel tagPAINT is then explored in cells co-expressing Halo-tagged CD3ζ and SNAP-tagged LAT in T-cells.

## 2. Methods and Materials

### 2.1 Plasmids

For the expression of Halo Tag-labelled human CD3ζ, the EGFP gene in the vector pEGFP-N1 was exchanged with the Halo Tag gene after restriction digest, generating an empty back bone where human CD3ζ could be inserted. For SNAP-CD3ζ the Halo tag was subsequently replaced with SNAP gene, using the AgeI and NotI sites. For SNAP-LAT, human CD3ζ of the SNAP-CD3ζ construct was replaced with human LAT

### 2.2 Synthesis of Halo tag-DNA ligands

5’amino modified DNA-docking strands were diluted in 10 mM sodium phosphate buffer pH 6.8, supplemented with 5 mM EDTA, to a final concentration of 1 mM. *N*-hydroxysuccinimidyl ester functionalised Halo tag ligand (NHS-HL) was freshly reconstituted in dry DMSO to a final concentration of 50 mM (unused NHS-HL was aliquoted and stored at −80°C). NHS-HL was diluted 10 times by adding 5’amino modified DNA-docking strands and mixing thoroughly. The reaction was left for 1 h at room temperature. The reaction product, HL functionalised with a DNA-docking strand was purified from excess unreacted NHS-HL by size exclusion chromatography, with 10 mM Tris supplemented with 1 mM EDTA as the mobile phase. Purified HL-DNA-docking strands were aliquoted and stored at −20°C until use (final concentration approximately 100-200 μM).

### 2.3 Synthesis of SNAP tag-DNA ligands

5’amino modified DNA-docking strands were diluted in 10 mM sodium phosphate buffer pH 6.8, supplemented with 5 mM EDTA, to a final concentration of 1 mM. *N*-hydroxysuccinimidyl ester functionalised SNAP ligand (O^6^-benzylguanine) was freshly reconstituted in dry DMSO to a final concentration of 50 mM (unused NHS-O^6^-benzylguanine was aliquoted and stored at −80°C). NHS-O^6^-benzylguanine was diluted 10 times by adding 5’amino modified DNA-docking strands and mixing thoroughly. The reaction was left for 1 hour at room temperature. The reaction product, O^6^-benzylguanine functionalised with a DNA-docking strand was purified from excess unreacted NHS-O^6^-benzylguanine by size exclusion chromatography, with 10 mM Tris supplemented with 1 mM EDTA as the mobile phase. Purified HL-DNA-docking strands were aliquoted and stored at −20°C until use (final concentration approximately 100-200 μM).

### 2.4 Transfection and fixation of Jurkat T-cells

E6.1 Jurkat T cells were transfected with Halo or SNAP constructs using an Invitrogen Neon Electroporation Transfection System (Life Technologies Pty Ltd.), using 3 pulses of 1350 V lasting 10 ms. The cells were left to recover after transfection in RPMI medium without Phenol Red (6040, GIBCO) supplemented 20 % (v/v) fetal bovine serum. Before seeding cells were pelleted by centrifugation, washed once with PBS and then resuspended in PBS. Cells we then used for seeding onto coverslips coated with 10 μg/mL anti-human CD3ε (activating) for 10 min at 37°C with 5% CO2, after which non-adherent cells were washed away with PBS and then fixed with freshly prepared warm 4% (w/v) PFA in PBS for 10 mins. Fixative was then washed away with PBS and then cells were permeabilised for labelling with 0.1% (v/v) Triton X-100 in PBS for 3 min.

### 2.4 Immunostaining for SNAP and Halo tags

Cells expressing Halo and SNAP tags were processed for staining in an identical manner. After fixation, cells were permeabilised with PBS supplemented with 0.2% Triton-X100 for 3 mins, and then subsequently incubated with 50 mg/mL BSA in PBS for 30 mins. After which the cells were stained with either anti-SNAP-ATTO488 or anti-Halo-ATTO568 (or both **Figure 2**) at a final concentration of 1 ug/mL in 50 mg/mL BSA in PBS for 30 mins. Cells were then washed with PBS and post-fixed using 4% PFA in PBS.

### 2.5 Labelling with SNAP and Halo DNA ligands

Both DNA-functionalised SNAP and Halo ligands were incubated with fixed and immunostained cell samples at a final concentration of 5 μM in PBS supplemented with 0.2% Tween-20 (PBST) for 10 min. The samples were then vigorously washed with 1 mL of PBST, several times to remove any non-specifically adsorbed ligand. Finally, the samples were incubated with gold nanorods (Nanopartz) in PBST for 10 mins before mounting for tagPAINT imaging.

### 2.6 tagPAINT imaging

Prior to imaging labelled cells, glass coverslips were mounted into a chamlide chamber and freshly prepared imaging strands (2nM ATTO655 P01 or 2nM ATTO655 P03) in PBST supplemented with 500 mM NaCl were added to the chamber. For tagPAINT imaging, ATTO655 imager binding was acquired with the 642 nm (0.075 kW/cm^2^) laser lines, respectively. For standard tagPAINT imaging an integration time of 80 ms was used, with a TIRF angle of 66.90°, with 40,000 frames acquired. Images were acquired as 512×512 pixel images with a pixel size of 97 nm. For exchange PAINT imaging (**Figure 2**) the same imaging parameters as above were used, but in between each image series the imager solution was exchanged by washing with at least 10 x 1mL PBST supplemented with 500 mM NaCl, and then adding in the second imager solution.

### 2.7 tagPAINT processing and image convolution

tagPAINT images were processed using Zeiss Zen Black software. The position of bound imaging strands in the acquisition was determined by Gaussian fitting, using a peak mask radius size of 6 pixels and a signal to noise ratio cut-off of 8. The localisation data was then drift corrected using the point patterns generated from the localisation of gold nanorod fiducials within the field of view using the Zeiss Zen Black software drift correction. The point pattern data was then convolved with a gaussian kernel using the ThuderSTORM plugin in Fiji, with a pixel size of 9.7 nm.

## 3. Results and Discussion

### 3.1 Specific imaging of SNAP and Halo-tagged proteins in knockout T-cell lines

To demonstrate the potential of this approach for DNA-PAINT we measured the specificity of the DNA-ligands and their targeting to the tagged proteins. To this end, we targeted SNAP and Halo tagged proteins in the immune synapse, LAT and CD3ζ (**Figure 1**). Halo and SNAP tagged proteins bind covalently to a single ligand, thus, functionalisation of these ligands with DNA target strands makes them amenable to DNA-PAINT imaging. Here we used 5’-amine functionalised DNA target strands and reacted them either with excess of O^6^-benzylguanine -NHS or Halo ligand-NHS ligands in a single step reaction (see **Materials and Methods**). Cells that were deficient in both LAT and CD3ζ were reconstituted with tagged versions of these proteins via transient transfection (**Figure 1**). To detect cells expressing the tag-bearing proteins we used immunostaining with antibodies raised to the tag (**Figure 1 A-D**), as the tags, and the proteins they are based on, are not endogenously expressed in mammalian cells. SNAP tagged LAT and CD3ζ expressing cells were detected by anti-SNAP-ATTO488 and subsequently imaged using DNA-PAINT (**Figure 1A and B**). DNA-PAINT imaging revealed the presence of punctate signal within the immune synapse, similar to that observed previously [27, 28](**Figure 1A and B**), which is absent when cell express the SNAP tag, but have not been labelled with the DNA-SNAP-ligand (**Figure S1A**). Peak localisation precisions were 12 nm (**Figure S2A**) and 10 nm (**Figure S2B**) for LAT and CD3ζ, respectively. For Halo tagged LAT and CD3ζ expressing cells were detected by anti-Halo-ATTO568, and again imaged by DNA-PAINT (Figure 1C and D). Again, similar to SNAP tag, punctate signal is observed again at the immune synapse, and is not seen in the absence of the ligand (**Figure S1B**). Peak localisation precision for Halo tagged (**Figure S2C and D**) for LAT and CD3ζ was 10 nm (5 nm contribution in CD3ζ was from a fiducial which could not be segmented from the data). Thus, labelling of tag for DNA-PAINT imaging appears to be highly specific.

**Figure 1.**
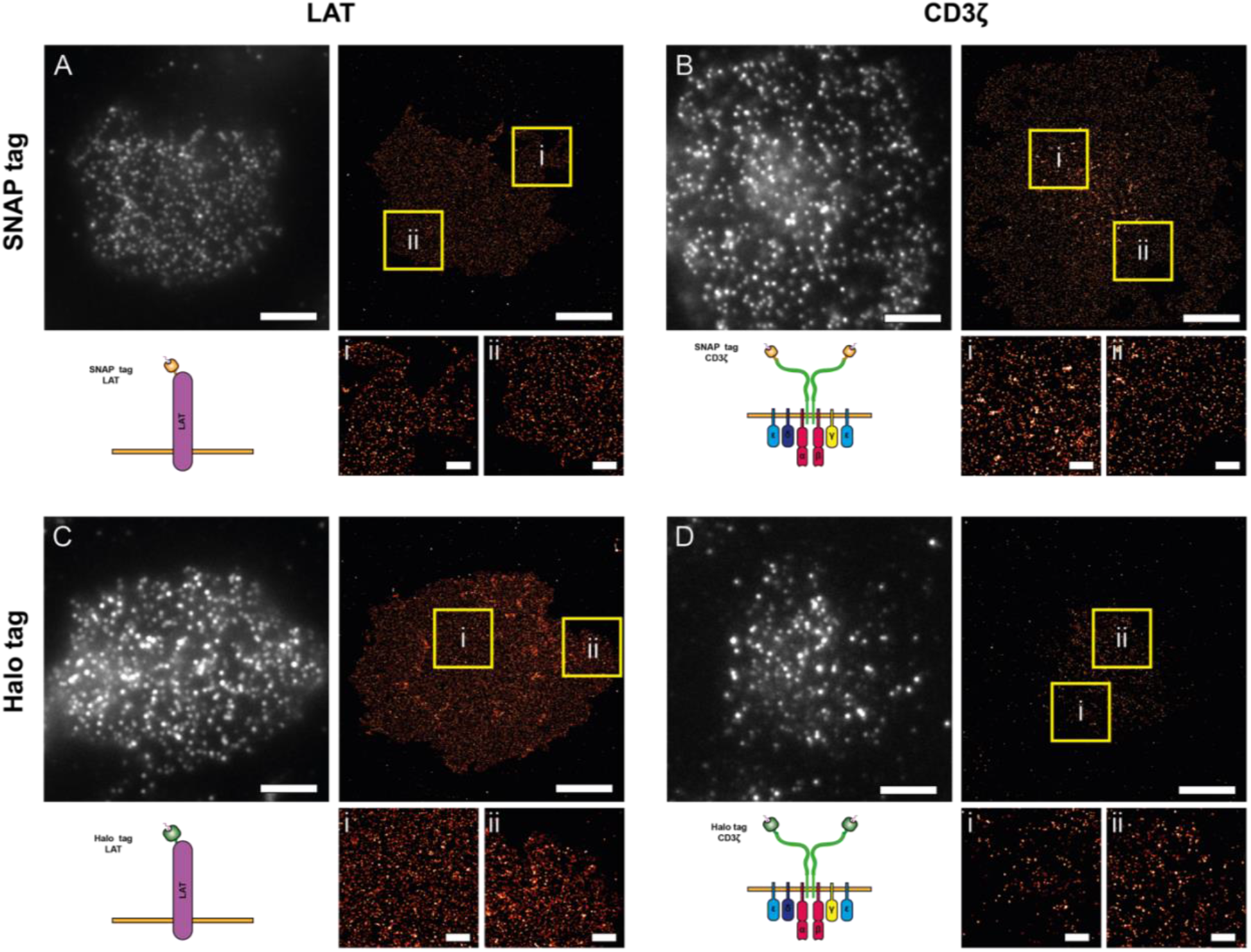
tagPAINT imaging of CD3ζ and LAT proteins. LAT and CD3ζ knock out (KO) Jurkat cells were transfected with LAT-SNAP **(A)**, CD3ζ-SNAP **(B)**, LAT-Halo **(C)**, CD3ζ-Halo **(D)** constructs. **A**) LAT-SNAP transfected cells were stained with anti-SNAP-ATTO488 antibody to determine expressing cells (top right). Gaussian convolved images of positive cells imaged using DNA-PAINT (top left; 2nM, P01-ATTO655 imager). Zoomed regions of DNA-PAINT imaging are shown (yellow boxes, i and ii). **B)** CD3ζ-SNAP transfected cells were stained with anti-SNAP-ATTO488 antibody to determine expressing cells (top right). Gaussian convolved images of positive cells imaged using DNA-PAINT (top left; 2 nM, P01-ATTO655 imager). Zoomed regions of DNA-PAINT imaging are shown (yellow boxes, i and ii). **C)** LAT-Halo transfected cells were stained with anti-Halo-ATTO568 antibody to determine expressing cells (top right). Gaussian convolved images of positive cells imaged using DNA-PAINT (top left; 2 nM, P03-ATTO655 imager). Zoomed regions of DNA-PAINT imaging are shown (yellow boxes, i and ii). **D)** CD3ζ-Halo transfected cells were stained with anti-Halo-ATTO568 antibody to determine expressing cells (top right). Gaussian convolved images of positive cells imaged using DNA-PAINT (top left; 2 nM, P03-ATTO655 imager). Zoomed regions (yellow boxes, i and ii). Scale bars for all larger images are 5 μm and for zoomed images 1μm.

### 3.2 Dual-tagPAINT imaging of CD3ζ-Halo and LAT-SNAP in the same cell

Having achieved specific targeting of the both SNAP and Halo-tagged proteins for DNA-PAINT in isolation, we sought to demonstrate the potential for imaging the two tags within the same cell using exchange DNA-PAINT. We used CD3ζ-knockout Jurkat T-cells and expressed both a Halo-tagged CD3ζ chain and SNAP tagged LAT protein (**Figure 2**). These proteins are involved in the early signalling of T-cell activation and have been shown to associate and co-cluster upon engagement of the TCR [29]. Here, each protein was labelled with a ligand bearing different DNA target sequences; P01 for SNAP-LAT and P03 for CD3ζ-Halo). Cells were stained with antibodies to the tags, as shown in **Figure 1**, to allow identification of cells expressing both tagged proteins (**Figure 2A and B**). First, LAT-SNAP was imaged by DNA-PAINT using P01 imager (**Figure 2C**), the sample was then washed with large volumes of imaging buffer, and then CD3ζ-Halo was imaged using the P03 imager (**Figure 2D**). Intermixing of the two proteins was observed at the immune synapse, as reported previously [29]. Thus, it is possible to orthogonally target and image different proteins within cells by employing the two tagging methods simultaneously.

**Figure 2.**
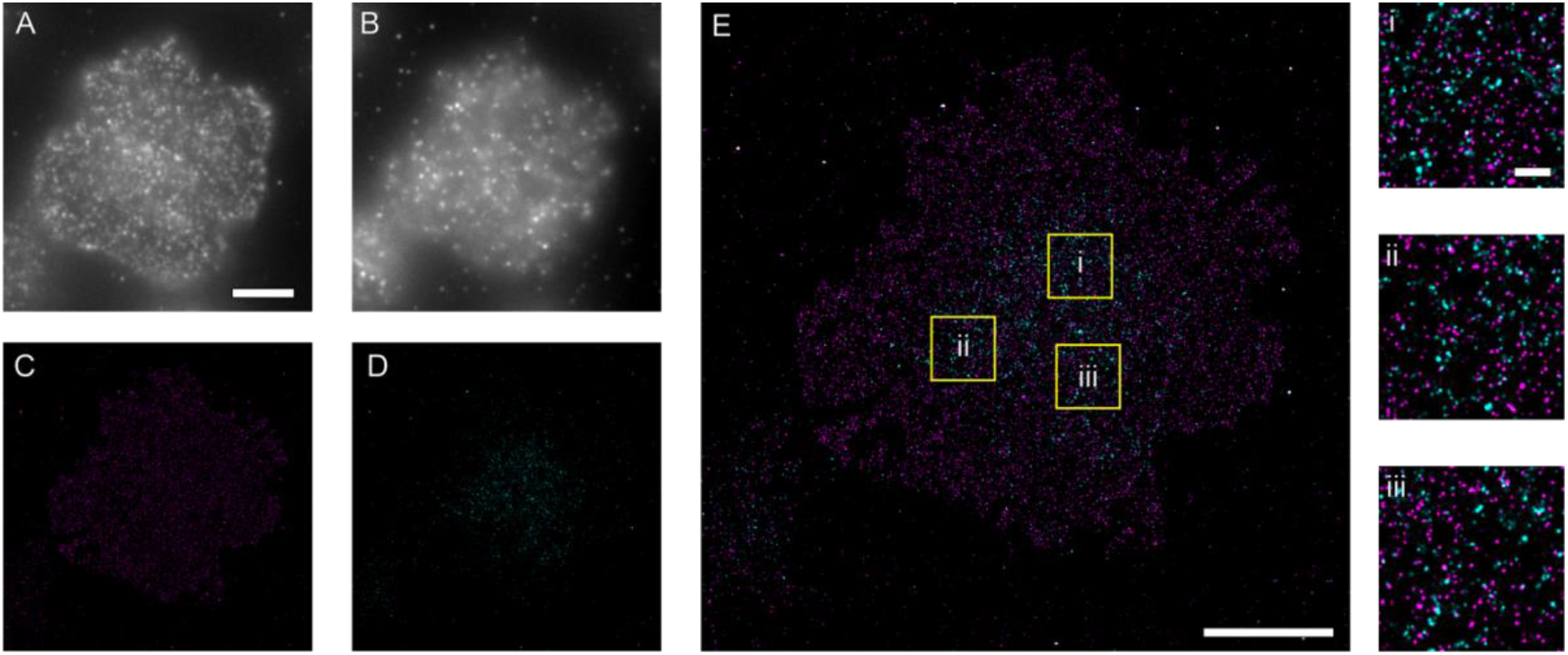
Dual tagPAINT imaging. CD3ζ knock out (KO) Jurkat cells were transfected with both CD3ζ-Halo and LAT-SNAP constructs. Cells were stained with both **(A)** anti-SNAP-ATTO488 and **(B)** anti-Halo-ATTO568 to identify cells expressing both constructs. Convolved images of Exchange PAINT imaging of **(C)** LAT-SNAP (2 nM, P01 ATTO655 imager) and **(D)** CD3ζ-Halo (2 nM, P01 ATTO655 imager), which are overlaid in E. Zoomed regions of the overlaid images are shown from regions i-iii (yellow boxes) in E (right).

## 4. Conclusion

Here we successfully conjugated DNA target strands to SNAP and Halo tag-labelled proteins. First, we demonstrated the specificity of the approach by targeting tagged CD3ζ and LAT expressed in knockout cells. Further, we showed the potential for dual colour imaging of these proteins (using both tags in the same cell). Given that the tags bind covalently only to a single ligand, and thus bear a single DNA target strand, this work provides a route to parallel quantitative DNA-PAINT imaging where multiple proteins are observed in this manner inside the same cell.

## SUPPLEMENTARY FIGURES

**Figure S1.**
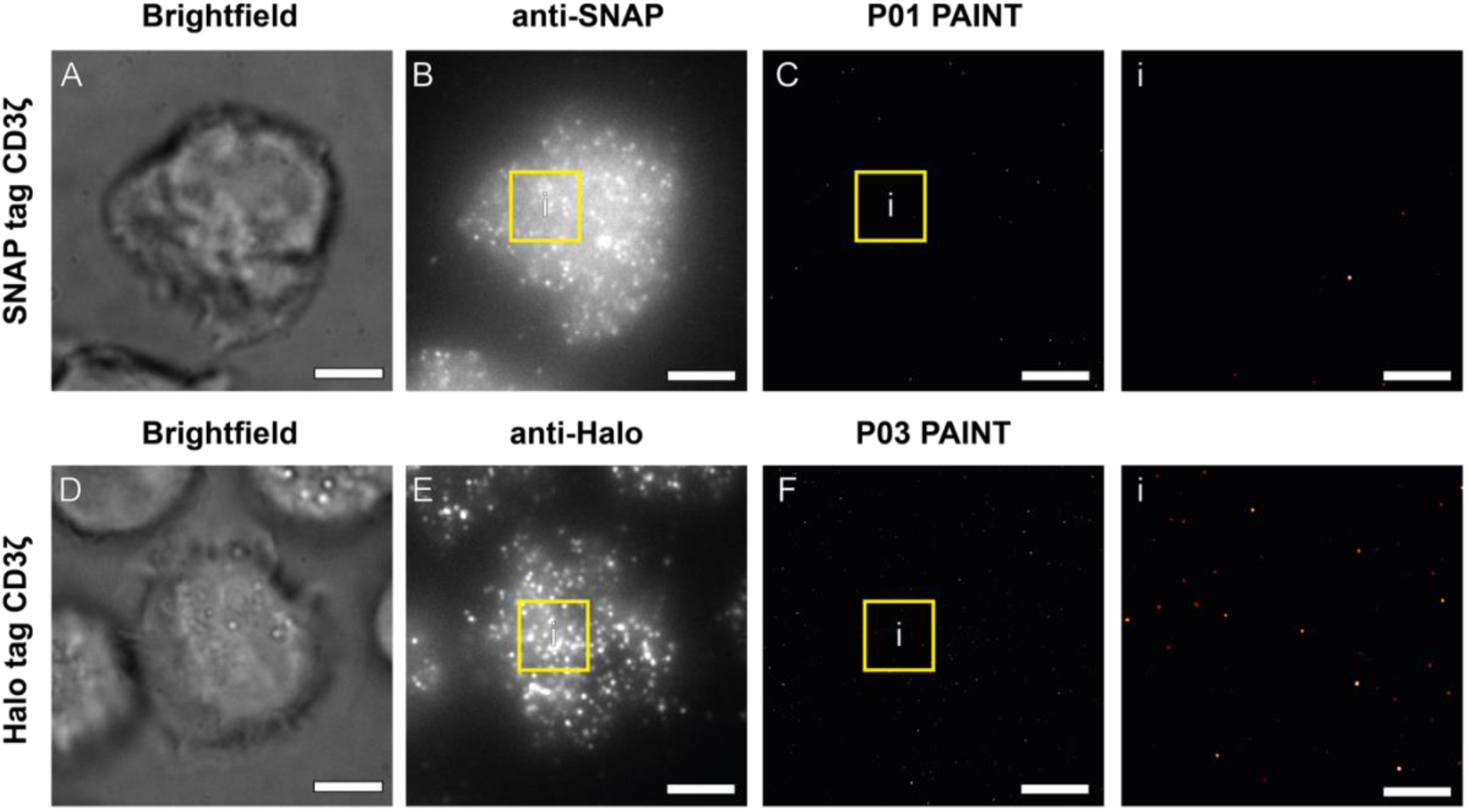
Background from tagPAINT imaging in the absence of tag ligand labelling. CD3ζ KO Jurkat cells expressing SNAP-CD3ζ **(A-C)** and Halo-CD3ζ **(D-F)** were processed and imaged using DNA-PAINT as in Figure 1, however, the ligand for the SNAP and Halo tags was not incubated with the cells prior to imaging. **A)** Brightfield of CD3ζ KO Jurkat cells expressing SNAP-CD3ζ. **B)** Anti-SNAP-ATTO488 staining. **C)** DNA-PAINT imaging (2nM, P01 ATTO655 imager), with zoomed region from the centre of the cell (i, yellow box) shown (right). **D)** Brightfield of CD3ζ KO Jurkat cells expressing Halo-CD3ζ. **B)** Anti-Halo-ATTO568 staining. **C)** DNA-PAINT imaging (2nM, P03 ATTO655 imager), with zoomed region from the centre of the cell (i, yellow box) shown (right). Scale bars for all larger images are 5 μm and for zoomed images 1μm.

**Figure S2.**
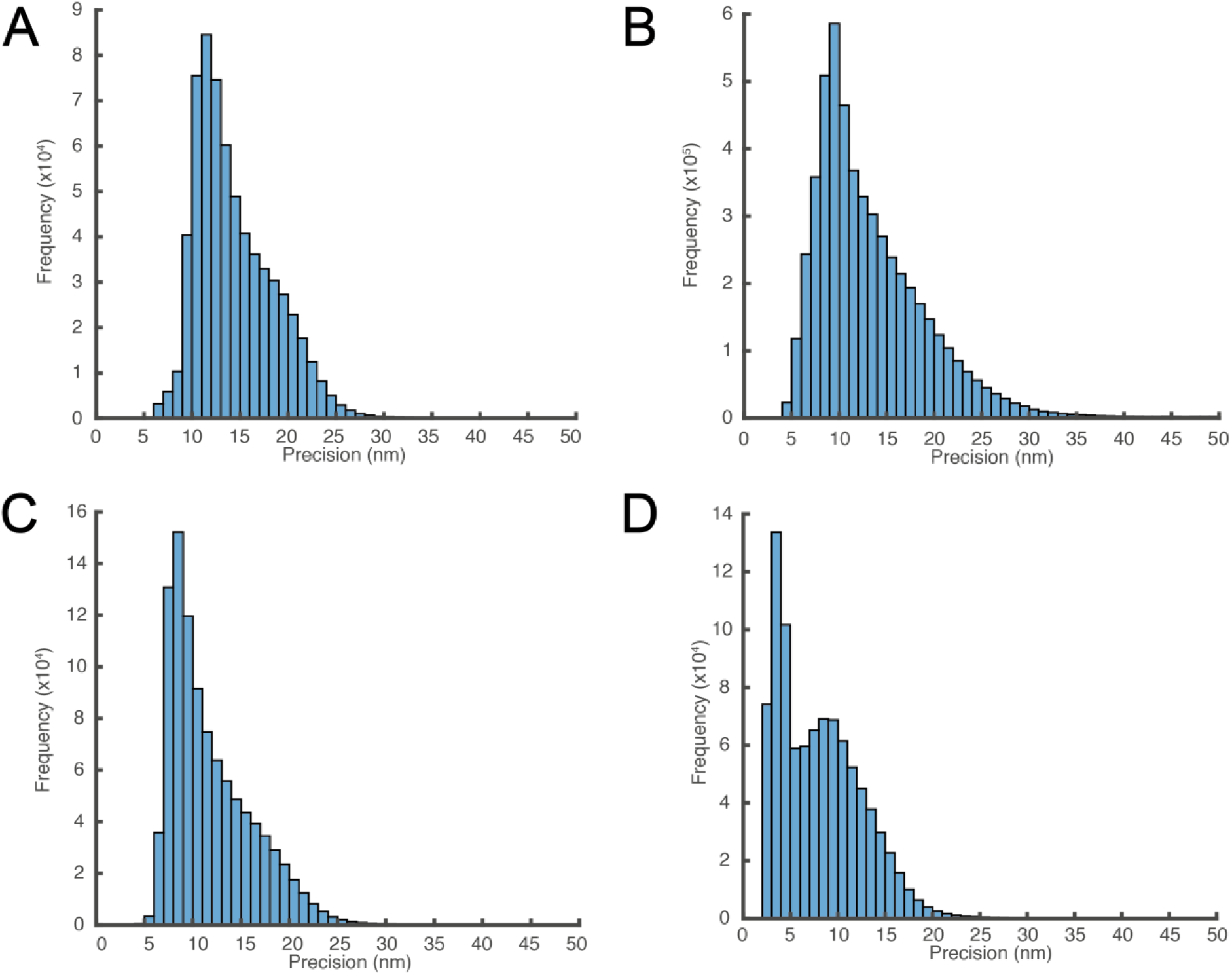
Localisation precision histrograms of tagPAINT imaging from Figure1. The localisation precision was calculated for each event within the image acquisition according to Mortensen *et al.* [30]. **A)** Localisation data from LAT-SNAP tagPAINT data (**Figure 1A). B)** Localisation data from CD3ζ-SNAP tagPAINT data (**Figure 1B). C)** Localisation data from LAT-Halo tagPAINT data (**Figure 1C). D)** Localisation data from CD3ζ-Halo tagPAINT data (**Figure 1D**).

## Author contributions

DJN conceived the study, designed the experiments, synthesised DNA tagging reagents, acquired data, wrote and drafted the manuscript, GH aided in experimental design, method optimisation and analysed data, JT aided in experimental design, imaging, and labelling/method optimisation, ZY cloned and generated constructs, HHM cloned and generated constructs, MABB aided in experimental design and drafting of the manuscript, JG aided in ligand synthesis optimisation and drafting of the manuscript, KG aided in experimental design and aided in writing and drafting of the manuscript. All authors gave final approval for publication.

## Ethics

This article does not present research with ethical considerations.

## Acknowledgments

We would like to thank the Mark Wainwright Analytical Centre and particularly the Biomedical Imaging Facility for additional access to microscopy facilities for the work.

## References

1 Ehmann, N., van de Linde, S., Alon, A., Ljaschenko, D., Keung, X. Z., Holm, T., Rings, A., DiAntonio, A., Hallermann, S., Ashery, U. et al. 2014 Quantitative super-resolution imaging of Bruchpilot distinguishes active zone states. Nat Commun. 5, 4650. (10.1038/ncomms5650)

2 Whelan, D. R., Bell, T. D. 2015 Image artifacts in single molecule localization microscopy: why optimization of sample preparation protocols matters. Sci Rep. 5, 7924. (10.1038/srep07924)

3 Heinze, K. G., Costantino, S., De Koninck, P., Wiseman, P. W. 2009 Beyond photobleaching, laser illumination unbinds fluorescent proteins. J Phys Chem B. 113, 5225–5233. (10.1021/jp8060152)

4 Baskin, J. M., Prescher, J. A., Laughlin, S. T., Agard, N. J., Chang, P. V., Miller, I. A., Lo, A., Codelli, J. A., Bertozzi, C. R. 2007 Copper-free click chemistry for dynamic in vivo imaging. Proc Natl Acad Sci USA. 104, 16793–16797. (10.1073/pnas.0707090104)

5 Saxon, E., Bertozzi, C. R. 2000 Cell surface engineering by a modified Staudinger reaction. Science. 287, 2007–2010.

6 Barlag, B., Beutel, O., Janning, D., Czarniak, F., Richter, C. P., Kommnick, C., Goser, V., Kurre, R., Fabiani, F., Erhardt, M. et al. 2016 Single molecule super-resolution imaging of proteins in living Salmonella enterica using self-labelling enzymes. Sci Rep. 6, 31601. (10.1038/srep31601)

7 Hauke, S., von Appen, A., Quidwai, T., Ries, J., Wombacher, R. 2017 Specific protein labeling with caged fluorophores for dual-color imaging and super-resolution microscopy in living cells. Chem Sci. 8, 559–566. (10.1039/c6sc02088g)

8 Nikic, I., Estrada Girona, G., Kang, J. H., Paci, G., Mikhaleva, S., Koehler, C., Shymanska, N. V., Ventura Santos, C., Spitz, D., Lemke, E. A. 2016 Debugging Eukaryotic Genetic Code Expansion for Site-Specific Click-PAINT Super-Resolution Microscopy. Angew Chem Int Ed Engl. 55, 16172–16176. (10.1002/anie.201608284)

9 Nikic-Spiegel, I. 2018 Genetic Code Expansion-and Click Chemistry-Based Site-Specific Protein Labeling for Intracellular DNA-PAINT Imaging. Methods Mol Biol. 1728, 279–295. (10.1007/978-1-4939-7574-7_18)

10 Wilmes, S., Staufenbiel, M., Lisse, D., Richter, C. P., Beutel, O., Busch, K. B., Hess, S. T., Piehler, J. 2012 Triple-color super-resolution imaging of live cells: resolving submicroscopic receptor organization in the plasma membrane. Angew Chem Int Ed Engl. 51, 4868–4871. (10.1002/anie.201200853)

11 Tyagi, S., Lemke, E. A. 2013 Genetically encoded click chemistry for single-molecule FRET of proteins. Methods Cell Biol. 113, 169–187. (10.1016/B978-0-12-407239-8.00009-4)

12 Bosch, P. J., Correa, I. R., Jr., Sonntag, M. H., Ibach, J., Brunsveld, L., Kanger, J. S., Subramaniam, V. 2014 Evaluation of fluorophores to label SNAP-tag fused proteins for multicolor single-molecule tracking microscopy in live cells. Biophys J. 107, 803–814. (10.1016/j.bpj.2014.06.040)

13 Agasti, S. S., Wang, Y., Schueder, F., Sukumar, A., Jungmann, R., Yin, P. 2017 DNA-barcoded labeling probes for highly multiplexed Exchange-PAINT imaging. Chem Sci. 8, 3080–3091. (10.1039/c6sc05420j)

14 Jungmann, R., Avendano, M. S., Woehrstein, J. B., Dai, M., Shih, W. M., Yin, P. 2014 Multiplexed 3D cellular super-resolution imaging with DNA-PAINT and Exchange-PAINT. Nat Methods. 11, 313–318. (10.1038/nmeth.2835)

15 Schnitzbauer, J., Strauss, M. T., Schlichthaerle, T., Schueder, F., Jungmann, R. 2017 Superresolution microscopy with DNA-PAINT. Nat Protoc. 12, 1198–1228. (10.1038/nprot.2017.024)

16 Nieves, D. J., Gaus, K., Baker, M. A. B. 2018 DNA-Based Super-Resolution Microscopy: DNA-PAINT. Genes (Basel). 9, (10.3390/genes9120621)

17 Jungmann, R., Avendano, M. S., Dai, M., Woehrstein, J. B., Agasti, S. S., Feiger, Z., Rodal, A., Yin, P. 2016 Quantitative super-resolution imaging with qPAINT. Nat Methods. 13, 439–442. (10.1038/nmeth.3804)

18 Schlichthaerle, T., Eklund, A. S., Schueder, F., Strauss, M. T., Tiede, C., Curd, A., Ries, J., Peckham, M., Tomlinson, D. C., Jungmann, R. 2018 Site-Specific Labeling of Affimers for DNA-PAINT Microscopy. Angew Chem Int Ed Engl. (10.1002/anie.201804020)

19 Sograte-Idrissi, S., Oleksiievets, N., Isbaner, S., Eggert-Martinez, M., Enderlein, J., Tsukanov, R., Opazo, F. 2019 Nanobody Detection of Standard Fluorescent Proteins Enables Multi-Target DNA-PAINT with High Resolution and Minimal Displacement Errors. Cells. 8, (10.3390/cells8010048)

20 Fabricius, V. L., J.; Geertsema, H.; Marino, S. F.; Ewers, H. 2018 Rapid and efficient C-terminal labeling of nanobodies for DNA-PAINT. Journal of Physics D: Applied Physics. 51,

21 Strauss, S., Nickels, P. C., Strauss, M. T., Jimenez Sabinina, V., Ellenberg, J., Carter, J. D., Gupta, S., Janjic, N., Jungmann, R. 2018 Modified aptamers enable quantitative sub-10-nm cellular DNA-PAINT imaging. Nat Methods. 15, 685–688. (10.1038/s41592-018-0105-0)

22 Gautier, A., Juillerat, A., Heinis, C., Correa, I. R., Jr., Kindermann, M., Beaufils, F., Johnsson, K. 2008 An engineered protein tag for multiprotein labeling in living cells. Chem Biol. 15, 128–136. (10.1016/j.chembiol.2008.01.007)

23 Los, G. V., Encell, L. P., McDougall, M. G., Hartzell, D. D., Karassina, N., Zimprich, C., Wood, M. G., Learish, R., Ohana, R. F., Urh, M. et al. 2008 HaloTag: a novel protein labeling technology for cell imaging and protein analysis. ACS Chem Biol. 3, 373–382. (10.1021/cb800025k)

24 Klein, T., Loschberger, A., Proppert, S., Wolter, S., van de Linde, S., Sauer, M. 2011 Live-cell dSTORM with SNAP-tag fusion proteins. Nat Methods. 8, 7–9. (10.1038/nmeth0111-7b)

25 Gu, G. J., Friedman, M., Jost, C., Johnsson, K., Kamali-Moghaddam, M., Pluckthun, A., Landegren, U., Soderberg, O. 2013 Protein tag-mediated conjugation of oligonucleotides to recombinant affinity binders for proximity ligation. N Biotechnol. 30, 144–152. (10.1016/j.nbt.2012.05.005)

26 Singh, V., Wang, S., Chan, K. M., Clark, S. A., Kool, E. T. 2013 Genetically encoded multispectral labeling of proteins with polyfluorophores on a DNA backbone. J Am Chem Soc. 135, 6184–6191. (10.1021/ja4004393)

27 Rossboth, B., Arnold, A. M., Ta, H., Platzer, R., Kellner, F., Huppa, J. B., Brameshuber, M., Baumgart, F., Schutz, G. J. 2018 TCRs are randomly distributed on the plasma membrane of resting antigen-experienced T cells. Nat Immunol. 19, 821–827. (10.1038/s41590-018-0162-7)

28 Williamson, D. J., Owen, D. M., Rossy, J., Magenau, A., Wehrmann, M., Gooding, J. J., Gaus, K. 2011 Pre-existing clusters of the adaptor Lat do not participate in early T cell signaling events. Nat Immunol. 12, 655–662. (10.1038/ni.2049)

29 Lillemeier, B. F., Mortelmaier, M. A., Forstner, M. B., Huppa, J. B., Groves, J. T., Davis, M. M. 2010 TCR and Lat are expressed on separate protein islands on T cell membranes and concatenate during activation. Nat Immunol. 11, 90–96. (10.1038/ni.1832)

30 Mortensen, K. I., Churchman, L. S., Spudich, J. A., Flyvbjerg, H. 2010 Optimized localization analysis for single-molecule tracking and super-resolution microscopy. Nat Methods. 7, 377–381. (10.1038/nmeth.1447)

